# Gene Expression Profiling Unveils the Temporal Dynamics of CIGB-300-Regulated Transcriptome in AML Cells

**DOI:** 10.1101/2023.03.06.530131

**Authors:** Dania Vázquez-Blomquist, Ailyn C. Ramón, Mauro Rosales, George V. Pérez, Ailenis Rosales, Daniel Palenzuela, Yasser Perera, Silvio E. Perea

## Abstract

Protein kinase CK2 activity is implicated in the pathogenesis of various hematological malignancies like Acute Myeloid Leukemia (AML) that remains challenging concerning treatment. Consequently, here we used Illumina HT-12 microarray gene RNA expression profiling to study the molecular events that might support the anti-leukemic effect of CIGB-300 peptide which targets both CK2 substrates and the CK2α catalytic subunits on HL-60 and OCI-AML3 cell lines. As a result, 185 and 812 genes appeared significantly modulated in HL-60 cells at 30 min and 3 h of incubation with CIGB-300 for p< 0.01 and FC>│1.5│, respectively; while 222 and 332 genes appeared modulated in OCI-AML3 cells. Importantly, functional enrichment analysis evidenced that genes and transcription factors related to apoptosis, cell cycle, leukocyte differentiation, signaling by cytokines/interleukins, and NF-kB, TNF signaling pathways were significantly represented in AML cells transcriptomic profiles. The influence of CIGB-300 on these biological processes and pathways is dependent on the cellular background, in first place, and treatment duration. Of note, the impact of the peptide on NF-kB signaling was corroborated by the quantification of selected NF-kB target genes, as well as the measurement of p50 binding activity and soluble TNF-α induction. Quantification of CSF1/M-CSF and CDKN1A/P21 by PCR supports peptide effects on differentiation and cell cycle. Overall, here we explore for the first time the temporal dynamics of the gene expression profile regulated by CIGB-300 and provide fresh molecular clues concerning the antineoplastic effect of CIGB-300 in two relevant AML backgrounds.

## 1. Introduction

Acute Myeloid Leukemia (AML) is a heterogeneous hematologic malignancy characterized by high proliferation and ≥ 20% of undifferentiated myeloid progenitor cells (blasts) in bone marrow or peripheral blood [1], representing the most common acute leukemia in adults [2].

AML, among the most aggressive and lethal types of cancer, is often characterized by resistance to standard chemotherapy and poor long-term outcomes, in part due to the chromosomal alterations and gene mutations frequently found in leukemic blasts [3]. Technological advances have led to a remarkable improvement in our understanding of cancer through the implementation of large-scale genomic, transcriptomic and proteomic analyses [4, 5]. Several studies have demonstrated the use of gene differential expressions as effective tools for risk stratification of AML patients [6–9] or drug sensitivity [10]. Microarray technology has contributed to a better classification of acute leukemias [8, 11–14]. The value of microarray technology was earlier demonstrated to classify acute leukemia in myeloid and lymphoid [15]. Later on, microarray permitted the recognition of molecular subtypes in ALL patients [16]. This technology has also been used in the diagnosis and prognosis of AML as well as in the study of mechanisms of pathogenesis and therapeutics action in this disease [17].

Despite the progress in understanding AML biology and the use of novel technologies to improve disease characterization, chemotherapy and hematopoietic stem cell transplant are still the principal treatment approach for AML. Therapies targeting recurrent genetic mutations have been also gaining importance but less explored have been targeting leukemia cells with no mutations [18]. FDA has recently approved several non-cytostatic agents for the treatment of the patient, targeting important pathways in AML [19]. Nevertheless, there is a need for novel agents to combine with standard chemotherapy to efficiently eliminate leukemic cells and improve the outcomes.

Protein kinase CK2 hyperactivity is implicated in the pathogenesis of several hematological malignancies; high levels of CK2 appeared as a common denominator in all hematologic neoplasms, suggesting that CK2 inhibition could represent an attractive molecular target in AML [20, 21]. Only two compounds, the ATP-competitive inhibitor CX-4945, and the synthetic-peptide CIGB-300 have advanced to a clinical setting [3, 22]. CIGB-300 is a peptide originally designed to block the CK2-mediated phosphorylation through binding to the phosphoacceptor domain in the substrates [23]. However, recent studies have demonstrated that this inhibitor can interact with the CK2α catalytic subunit and regulate part of the CK2-dependent phosphoproteome in AML cell lines [21]. Additionally, proteomic analysis supported previous results evidencing that the pro-apoptotic effect, the impact over the cell cycle, the redox regulation, and the modulation of transcriptional/ translational processes are common denominators for CIGB-300-mediated CK2 inhibition in AML cells [21, 24]. However, a comprehensive characterization of the gene profile on CIGB-300-treated AML cells helping to understand its antileukemic effect, has not been accomplished yet.

Using microarray approach, here we interrogated the temporal gene expression profile modulated for CIGB-300 on HL-60 and OCI-AML3 cell lines uncovering key molecular events that might support the antileukemic effect of this CK2 peptide inhibitor.

## 2. Materials and Methods

### Cell Culture

Human AML cell lines HL-60 and OCI-AML3 were originally obtained from the American Type Culture Collection (ATCC, VA, USA) and the German Collection of Microorganisms and Cell Cultures (DSMZ, Braunschweig, Germany), respectively. Both cell lines were cultured in RPMI 1640 medium (Invitrogen, CA, USA) supplemented with 10% (v/v) fetal bovine serum (FBS, Invitrogen, CA, USA) and 50 μg/mL gentamicin (Sigma, MO, USA) under standard cell culture conditions at 37 °C and 5% CO_2_.

### Experiment setting up

An experiment with three replicates per condition of both cell lines was designed. The eight groups included untreated cells at 30min and 3h and treatment with 40μM of CIGB-300 peptide for 30min and 3h. After incubations cells were pickup in buffer RLT with 1% of β-mercaptoethanol and total RNA purification proceeded following the instructions of RNeasy Plus mini kit (Qiagen, USA). Quality control of total RNA was carried out by spectrophotometric readings of optic density (OD) at 260 and 280nm in Nanodrop 1000 (ThermoFisher, USA) to determine the concentration (> 80ng/µL) and OD260/280 ratio (1.8-2.2). Additionally, RIN (7-10) was calculated by capillary electrophoresis in a Bioanalyzer (Agilent, Waldbronn, Germany).

### Gene expression profile by Microarray

2.5 µg (100 ng/µL) of each total RNA sample was sent to McGill University and Génome Québec Innovation Centre (Montréal, Québec, Canada) for the experiment in Affymetrix Clariom S microarray gene expression platform.

### Quantitative PCR amplification

We obtained cDNA in 20 µL, from 870 ng of total RNA of three samples per group, following the instructions of the manufacturer of Transcriptor First Strand cDNA Synthesis Kit (Roche, Germany). qPCR reactions were set up in 20 µL with 300 nM of oligonucleotides (Table S1) and LightCycler® 480 SYBR Green I Master 2x (Roche, Germany) using three technical replicates per sample. The runs were carried out in a LightCycler®480II (Roche, Germany) equipment using the standard program SYBR Green Probe II and controls [25]. Ct and efficiency values were obtained and used in REST 2009[26] to report a Change Factor in gene levels after the treatment for 30min and 3h with CIGB-300, in relation to untreated cells after the normalization with GAPDH, DDX5, and ABL1 as reference genes [25]. The program reports a p-value after a Pair Wise Fixed Reallocation Randomization Test [27], reporting increasing and decreasing gene levels as UP and DOWN, respectively.

### Basic Microarray Data Analysis provided by the Genome Québec Innovation Centre

Basic bioinformatics analysis of microarray experiment was performed as a custom service at McGill University and Génome Québec Innovation Centre (Montréal, Canada). This service included quality control, preprocessing, and exploratory and differential expression analysis. As a result of the quality control, the array corresponding to replicate 1 of the OCI-AML3 control cell at 30min (O-30minC-1) was removed from the analysis. Preprocessing included the Robust Multi-array Average (RMA) method [28] that performs background adjustment using the RMA convolution model, followed by quantile normalization and log2 transformation. Probes belonging to the same gene are then averaged using a robust model that estimates probe-specific effects using all arrays. Technical replicate arrays were averaged within the groups defined by variable(s) SampleID. The exploratory analysis applied clustering and dimensionality reduction techniques to the expression profiles in a hypothesis-free manner such as hierarchical clustering based on the correlation distance, 2D Multidimensional scaling plot (MDS), and ANOVA R^2^ of covariates on the first Principal Components.

For differential expression analysis, the Bioconductor Limma package was used [29]. Statistical tests contrasting different treatments were performed (Moderated t-tests)[30]. The Benjamini-Hochberg was used for FDR estimation [31].

### Additional Bioinformatics Analysis

Genes with a fold change (FC) greater than [1.5] and p values lesser than 0.01 (p< 0.01), in each cell line at each time compared with the untreated control, were considered Differentially Expressed Genes (DEGs) and used for later bioinformatics analysis. Venn diagram was constructed to show common and different genes (VIB /UGent Bioinformatics & Evolutionary Genomics Gent, Belgic; http://bioinformatics.psb.ugent.be/webtools/Venn/).

Clustergrammer online tool was used to generate unsupervised heatmaps from the most differentially expressed 1100 genes [32]. Enrichr online analysis associated with Clustergrammer was used for enrichment exploration of the top 100 genes and other lists [33]. Additional functional enrichment analyses were carried out with the bioinformatic tool Metascape [34]. Metascape gene annotation and analysis resource (https://metascape.org/), is a web-based tool that computes the accumulative hypergeometric distribution and enrichment factors to identify significantly enriched biological processes through statistical analysis (p-value<0.01, enrichment factor >1.5)).

BisoGenet (version 3.0.0) Cytoscape plugin available from Cytoscape Application Manager, was used to generate PPI networks [36]. Cytoscape software (v.3.3.0) was used as a network framework [37]. Analysis on-networks to predict transcription factors (TF) from DEGs was carried out with the iRegulon plugin (version 1.3) [38]. Highly interconnected genes on the network were explored with the MCODE plugin (version 1.4.1) [39].

### P50-DNA binding activity detection by ELISA

To measure p50-DNA binding activity HL-60 and OCI-AML3 cells were incubated with CIGB-300 at a dose of 40 µM for 30 min, 2 h, and 5 h at 37 °C in 5% CO_2_. After treatment, cells were collected and nuclear extraction was performed using a Panomics kit (#EK110). The DNA-binding activity of the p50 NF-kB subunit was monitored using a commercially available ELISA-based assay (Panomics, #EK1111). Briefly, nuclear samples (10 µg) were incubated in an ELISA plate that was coated with oligonucleotides containing a p50 consensus regulatory element sequence. The positive control nuclear extract was prepared from HEK293 cells that were treated with 20ng/mL of TNF-α for 30 min. The wells were washed and exposed to a primary antibody specific for the p50 subunit of NF-kB. The binding of the primary antibody to protein was detected through a chromogenic reaction involving the enzymatic breakdown of 3, 3’, 5, 5’ tetramethylbenzidine via a horseradish peroxidase (HRP)-conjugate secondary antibody. Finally, absorbance at 450nm was read using a CLARIOstar® high-performance monochromator multimode microplate reader (BMG LABTECH, Ortenberg, Germany).

### TNF-α secretion detection by ELISA

To evaluate TNF-α secretion we employed a commercial human TNF-α ELISA kit (R&D Systems). HL-60 and OCI-AML3 cells were seeded at a concentration of 500000 cells /mL in 12-well cell culture plates and treated with CIGB-300 (40 µM) for 3 h and 24 h at 37 °C in 5% CO_2_. Phorbol myristate acetate (PMA) at a dose of 2.5 ng/mL was used as a positive control of TNF-α induction for 24 h of treatment. After incubation, supernatants were collected and subjected to ELISA analysis.

## 3. Results

### 3.1 Profiling of CIGB-300-regulated transcriptome in AML cells

Firstly, different diagnostic methods were used to sense microarray results (**Figure S1**). A hierarchical clustering based on the correlation distance is shown in **Figure S1A**. This one-dimensional clustering separates samples into experimental groups firstly based on cell line and later based on the treatment time and treatment received. Replicate 1 of the sample from OCI-AML3 at 30min (O-30min-1) was excluded from the analysis as it is an outlier. The analysis of ANOVA R^2^ of covariates on the first Principal Components showed main differences in the first components (PC1, PC2) come from cell lines differences; treatment (in PC3) and time (30min or 3h, in PC4) govern the differences in the other components (**Figure S1B**). The separation among the eight groups of this experiment in a homogeneous way is shown through Multidimensional Scaling plotting (**Figure S1C**). All these methods show the homogeneity of the data per group, the proper separation among them, and the main and ranked sources of variabilities of this microarray experiment.

The number of Differentially Expressed Genes (DEGs) obtained in each comparison, for a FC >[1.5] and a p< 0.01 are shown in **Figure 1A**. As a result, in HL-60 cells, 185 and 812 genes were identified as significantly modulated at 30 min and 3 h, while in OCI-AML3 cells, 222 and 332 genes appeared differentially modulated in response to CIGB-300 treatment. Interestingly, the numbers of genes up-regulated are always higher than down-regulated. HL-60 showed a much higher number of regulated genes after 3h of treatment.

**Figure 1.**
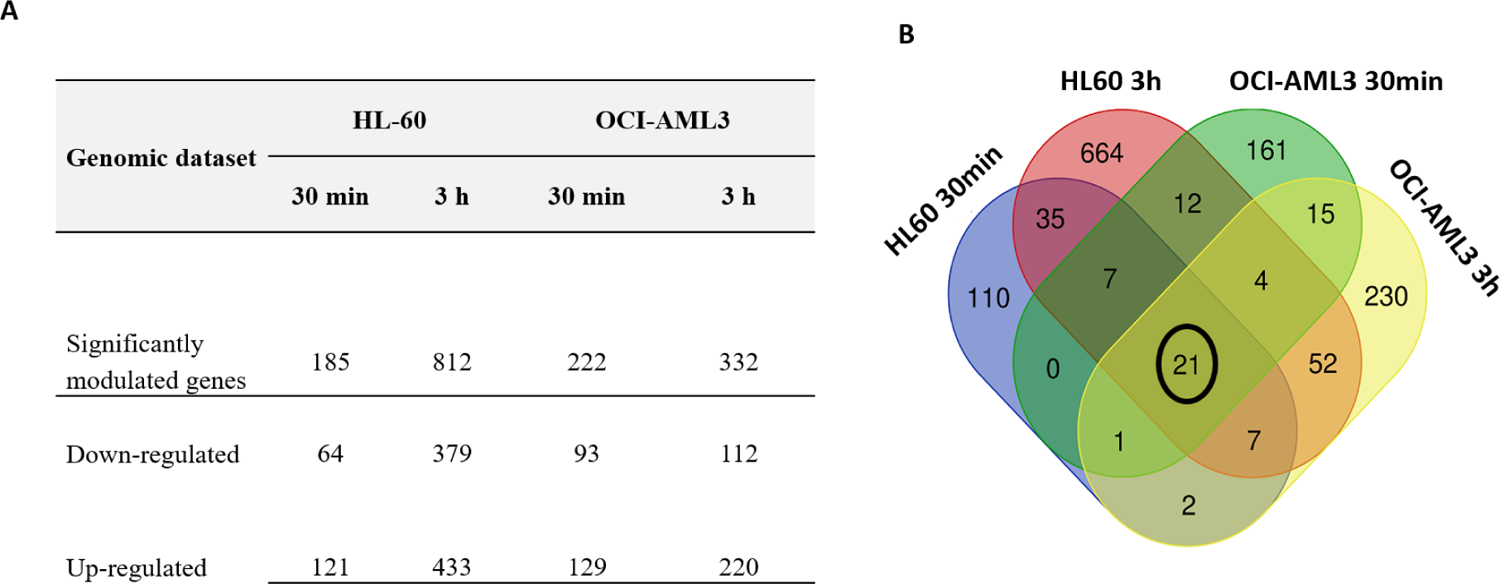
Genomic profile and Differentially Expressed Genes (DEGs) of AML cells treated with CIGB-300 peptide. (A) Numbers of differentially modulated genes in each AML cell line and time points; (B) Venn diagram of sets of DEGs for the treatment groups (p< 0.01;|FC|>=1.5); groups are represented in colors as HL-60 30min and 3h, OCI-AML3 30min and 3h. Numbers refer to DEGs specific to each treatment or common to two or more treatments. Inside the circle common, DEGs for the four groups.

Subsequently, a Venn diagram was constructed to show common and different genes (**Figure 1B**). Only 21 genes are shared in the four comparison groups, most of them are transcription factors (EGR1/2/3, FOS, FOSB, IER3, JUN, JUNB, NR4A1/2/3, TNF, ZFP36) or chemokines CCL2, CCL3 and CXCL8/IL8. It points to the recruitment of a rapid response to stress and inflammation to deal with the peptide effect.

An unsupervised Heatmap from the most differentially expressed 1100 genes using the online tool Clustergrammer showed very well-delimited clusters (**Figure 2A**). On top, there is a Cluster containing genes that increased their expression in both cell lines and time points (Cluster I) although with different kinetics and magnitudes. Heatmap of the top 100 up-regulated genes in Cluster I showed those common genes, biological pathways, and processes (**Figures S2A and B**). Even when the changes are more pronounced in HL-60 after 3h of treatments, it is interesting to note among the GO terms those related to cytokine-mediated signaling pathway, cellular response to cytokine stimulus, and inflammatory response sharing genes as CXCL10, TFN, IL1B, and other transcription factors, also included in positive regulation of transcription, and chemokines that participate in the Regulation of inflammatory response (**Figure 2B**). The main pathways are related to TNF and IL-17, TLR, NF-kB signaling, Cytokine-cytokine receptor interaction, and C-type lectin receptor signaling pathways (**Figure S2B**).

**Figure 2.**
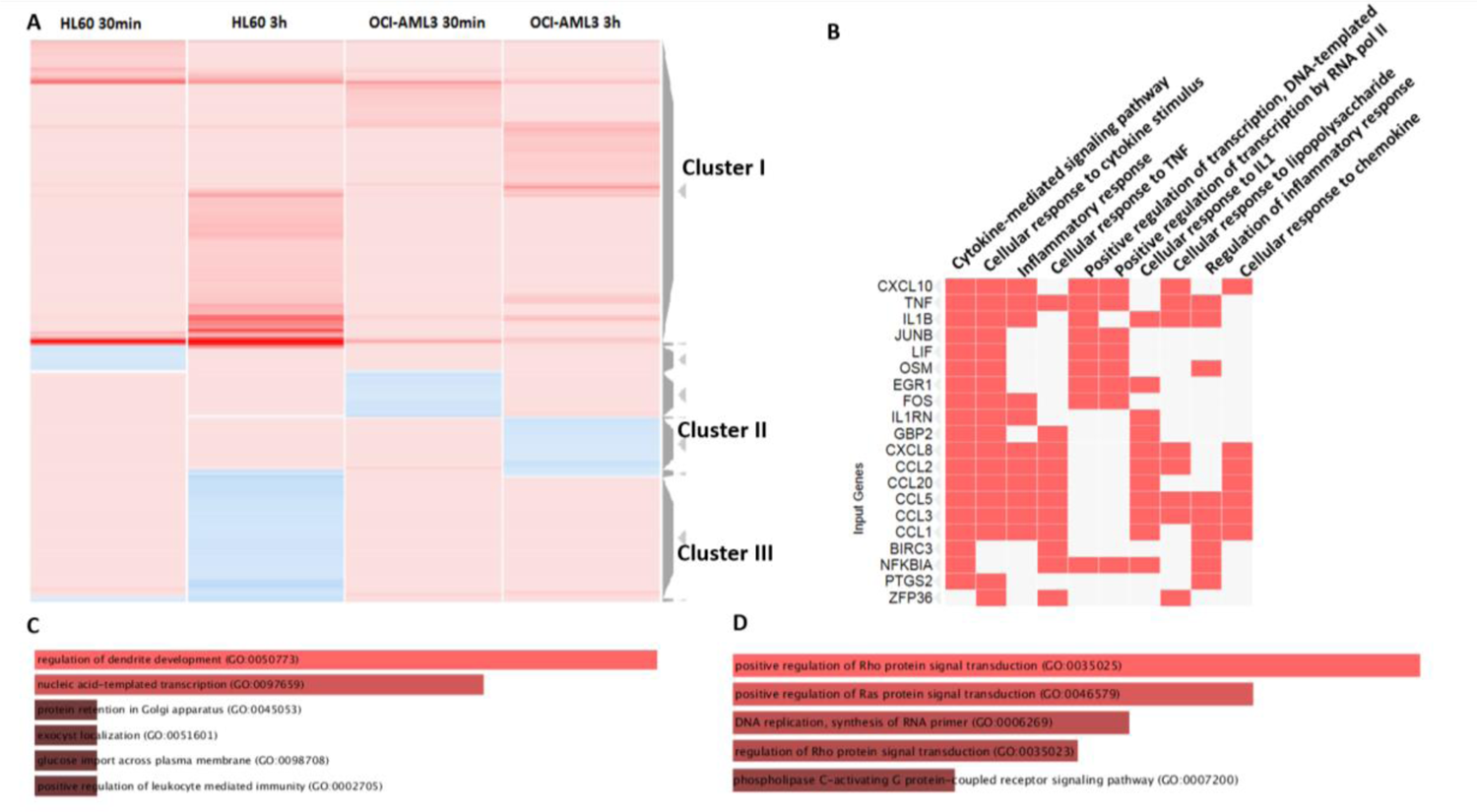
Analysis of DEGs by the action of CIGB-300 treatments and enriched biological processes. (A) Unsupervised Heatmap from the most differentially expressed 1100 genes, ranking the row order by Clusters. A subset with highly up-regulated genes (Cluster I), a subset where genes decreased their expression in OCI-AML3 at 3h but increased in HL-60 (Cluster II) and a subset where genes decreased their expression in HL-60 at 3h but increased in OCI-AML3 (Cluster III) are shown. (B) GO terms enriched using associated Enrichr analysis and genes included in more represented processes from Top 100 analysis in Cluster I, in a Clustergrammer shape. (C) and (D) represent the Enrichr analysis from genes included in Cluster II and III, respectively.

Clusters II and III show interesting behaviors related to cell lines background (**Figure 2A**), where genes decreased their expression in HL-60 at 3h but increased in OCI-AML3 and *vice versa*. Enrichment analysis using the Enrichr tool associated with Clustergrammer showed different biological processes containing those genes. While 3h after peptide treatment of HL-60 (**Figure 2D**), CIGB-300 decreased the expression of 232 genes mostly related to signaling transduction through Ras and Rho proteins and phospholipase C-activating G protein-coupled receptor, the 98 genes that decreased expression in OCI-AML3 (**Figure 2C**) are mainly related to other biological processes as regulation of dendrite development, transcription, protein retention in Golgi, glucose import or leukocyte mediated immunity regulation.

### 3.2 Enrichment analysis of DEGs pointed to cell line and timing dependence

When enriched biological processes were analyzed on each cell line per time of treatment, we obtained the Metascape pictures shown in **Figure 3**. Since 30 min of treatment in HL-60, there were genes from the NGF-stimulated transcription process, including transcription factors (EGR1/2/3/4, FOS, FOSB, JUN, SRF, NR4A1/2/3), and chemokines CCL2 & CXCL8. In both cell lines, the PID NFAT Transcription factor pathway is enriched which included similar 14 TF and chemokines genes together with TNF and additional 18 and 37 genes in HL-60 and OCI-AML3, respectively. Regulation of hemopoiesis and differentiation are also represented in both cell lines. The first process appeared for HL-60 (30 min and 3h) and OCI-AML3 (30 min); the second one was for HL-60 30 min and OCI-AML3 3h. The orexin receptor pathway is enriched in OCI-AML3 at 30 min and 3h, but only later at 3h in HL-60.

**Figure 3.**
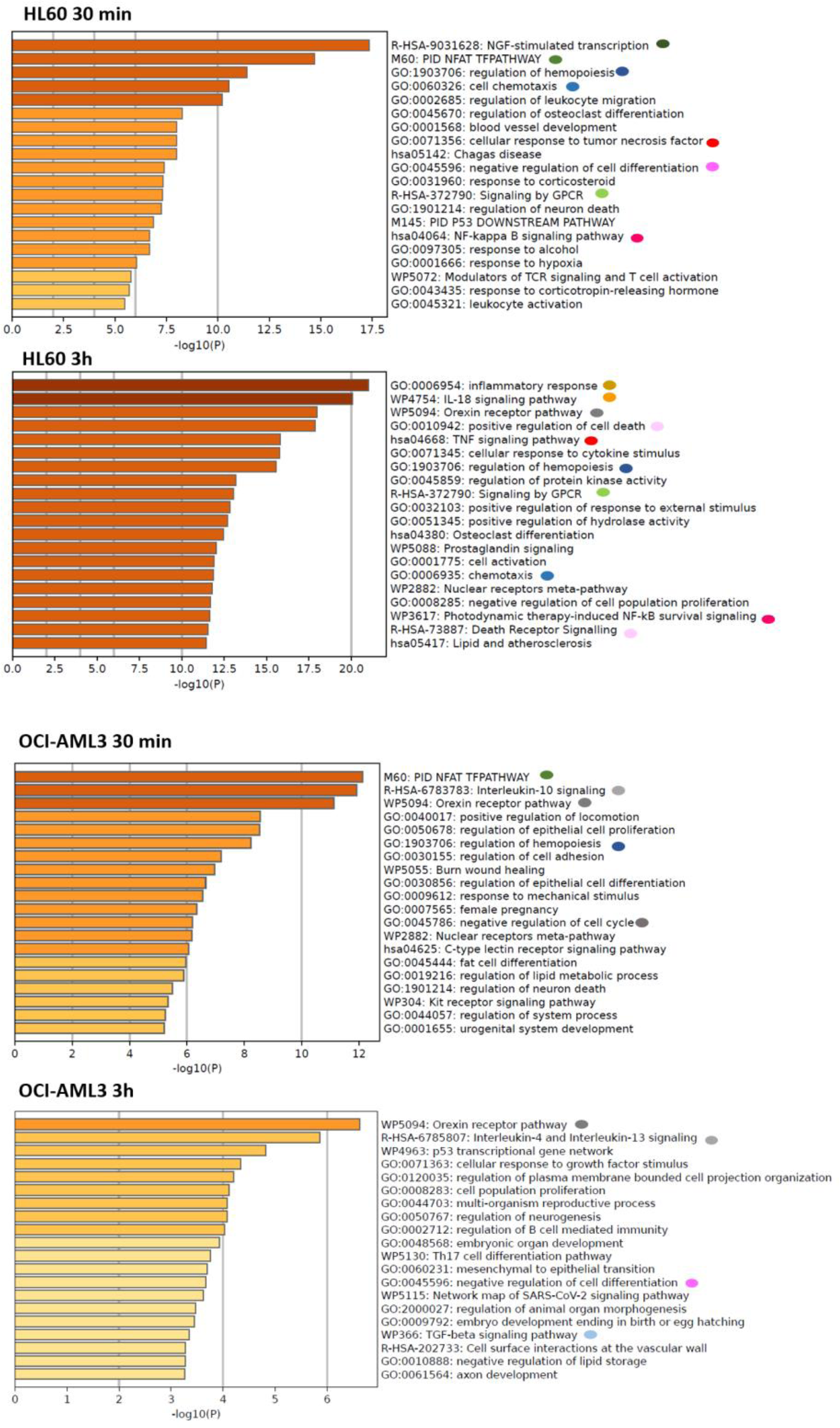
Metascape GO analysis for AML cell lines at 30min and 3h. The most enriched (-log10(P)) biological processes in HL-60 or OCI-AML3 after CIGB-300 treatment for 30min and 3h is shown. Colored circles represent GO terms of relevance; coincidences are marked with the same colors.

Of note, the cellular response to TNF, NF-kB signaling pathway and Signaling by GPCR are processes highly enriched in HL-60 treated for 30min with CIGB-300. After 3h of treatment the inflammatory response, IL-18 signaling pathway, and chemotaxis appeared as highly enriched processes in HL-60, together with TNF signaling pathway, signaling by GPCR regulation of protein kinase activity and positive regulation of cell death or Death Receptor Signaling. These last processes included genes as BCL2L11, CDKN1A, FAS, GADD45B, MAP3K5, SQSTM1 and TNFRSF12A.

In contrast, in OCI-AML3, IL10 and IL4/IL13 signaling appeared enriched at 30 min and 3h, respectively. At the earlier time, it showed the negative regulation of the cell cycle as an enhanced process.

### 3.3 DEGs show high functional network connections

Using Cytoscape as a framework, we investigated the network connection among genes in each cell line, also considering their expression kinetics (**Figure 4**).

**Figure 4.**
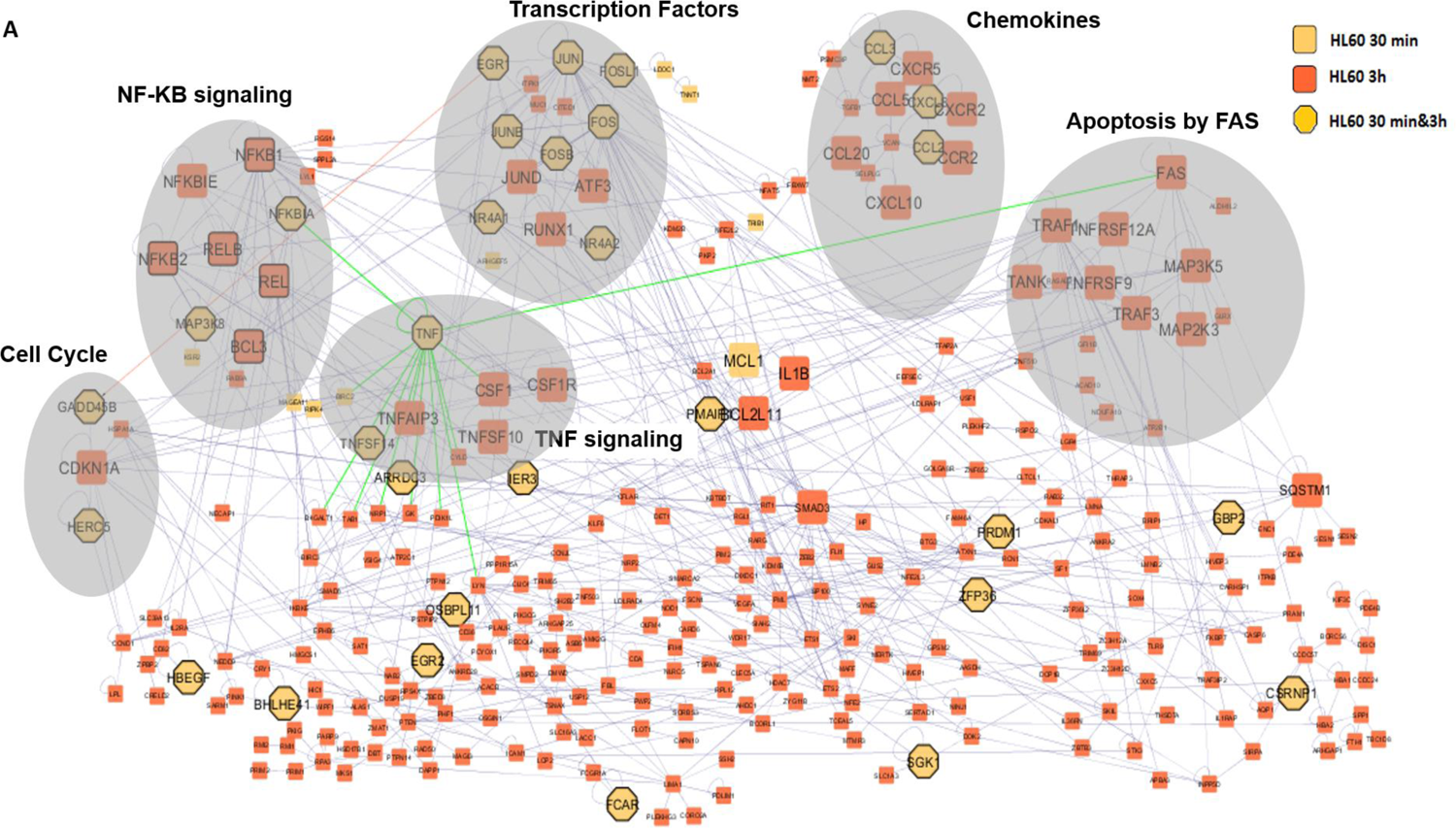
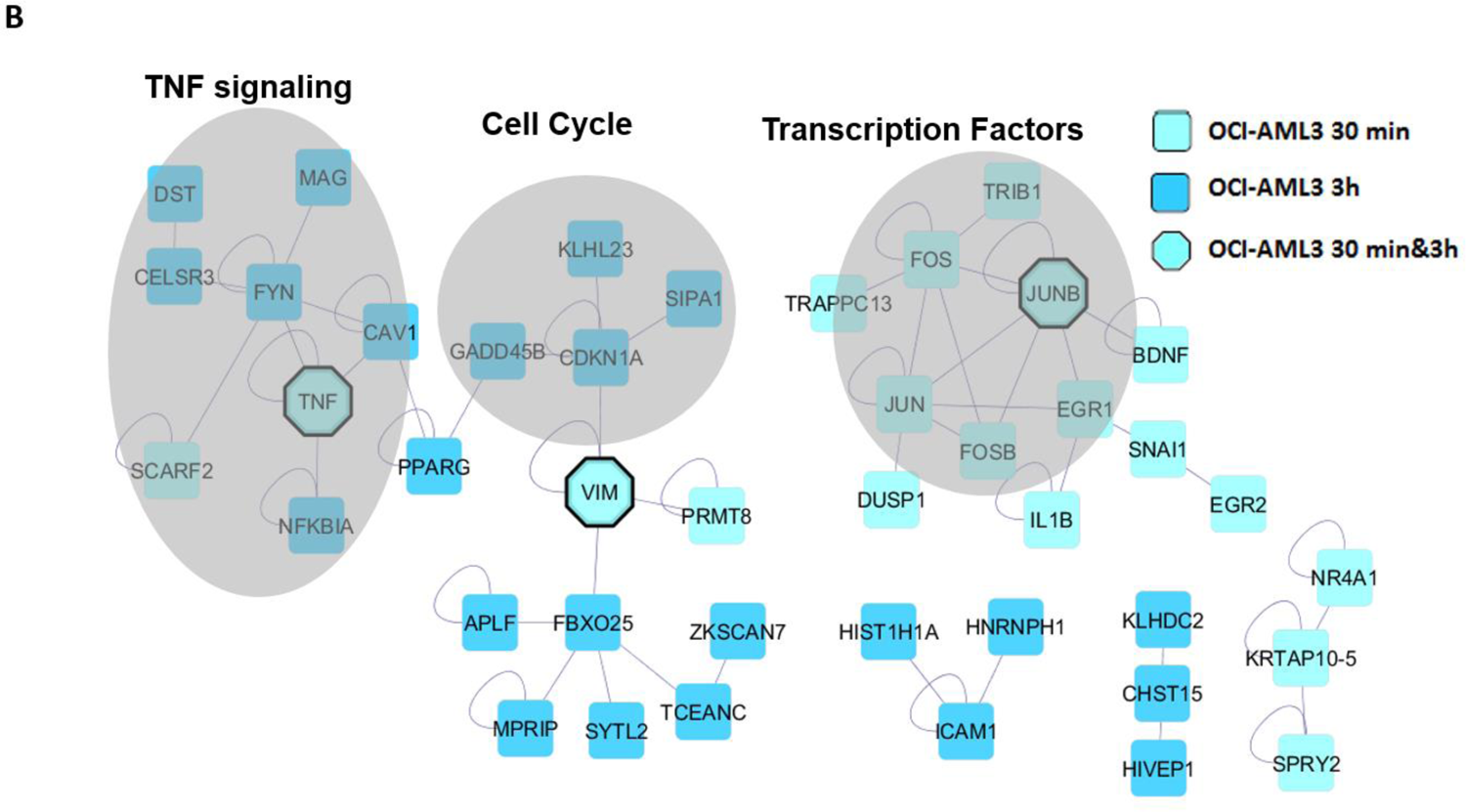
Network of DEGs by CIGB-300 treatment. Connected DEGs are shown in HL-60 (A) and OCI-AML3 (B) in both treatment times. Some modules of interest obtained by MCODE are shown in grey ovals. Additional TF are black lines surrounded in the net, as well as green edges of interest from TNF. The network was generated by the BisoGenet Cytoscape application.

In HL-60 cells, 305 out of 915 genes were connected in functional networks (**Figure 4A**). Most of the genes were regulated after 3h of treatment and 30 of them were regulated at both time points. The majority of these genes regulated along the experiment were transcription factors; additional TF highly connected are black lines surrounded in the net. The most interconnected modules included TF as EGR1, JUN, JUNB, FOS, FOSB, NF-kB family members, RUNX1, TNF, and genes connected to signaling for apoptosis and cell cycle control, such as FAS and CDKN1A. On the other side, DEGs in OCI-AML3 were less connected in networks. **Figure 4B** shows the main modules obtained with 42 out of 431 genes. Here, we also found highly connection among TF EGR1, JUN, JUNB, FOS, and FOSB but mainly at 30 min post-CIGB-300 treatment. TNF, regulated along all the experiment, connect with NF-KB regulator NFKBIA and with genes connected to signaling for apoptosis and cell cycle control as PPARG, GADD45B, and CDKN1A (p21). P21 is also connected to Vimentin (VIM), which is also regulated throughout the experiment.

### 3.4 Transcription factor prediction from DEGs shows differential dynamics in both AML cell lines

For a deeper understanding of gene expression regulation in HL-60 and OCI-AML3 after CIGB-300 treatment for 30 min and 3h, we explored gene regulatory motifs from DEGs networks using the iRegulon application. In **Figure 5** we show two graphs with ordered TF according to the Normalized Enrichment Score (NES) per cell line and time point. A high NES score (≥3.0) indicates a motif that recovers a large proportion of the input genes within the top of its ranking.

**Figure 5.**
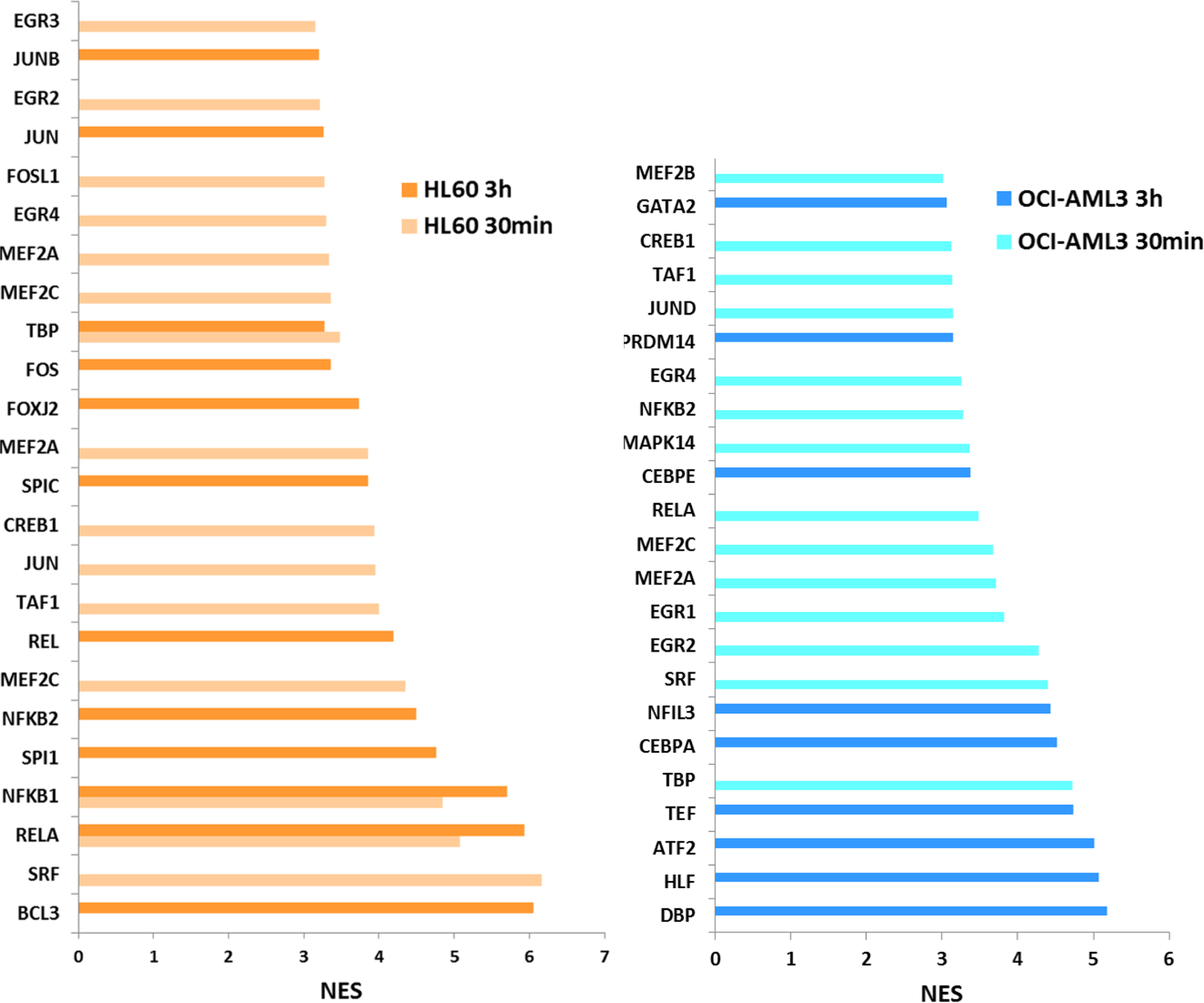
Predicted Transcription factors from DEGs. TF predicted from regulatory motifs in DEGs in HL-60 and OCI-AML3 after 30 min and 3h of CIGB-300 treatment are shown in both panels of the Figure. The Cytoscape on-net analysis plugin iRegulon was used and a NES>3.0 was selected.

In HL-60, gene transcription regulation is governed at 30 min by SRF, NF-kB family members (RELA, NFKB1), JUN, CREB1, MEF2A/C, TBP, and EGR 4/2/3; later at 3h, BCL3 gains importance together with NF-kB family members (RELA, NFKB1 and 2, REL), FOS, TBP, JUN, and JUNB. Otherwise, in OCI-AML3 the 30 min gene regulation is mainly accomplished by TBP, SRF, EGR2/1/4, MEF2A/C/B, RELA, NFKB2, MAPK14, JUND, and CREB1 while at 3h post-treatment the gene regulation strategy completely changed with a role for DBP, HLF, ATF2, TEF, CEBPA/B, NFIL3, and GATA2 and not by NF-kB family members or AP-1 complex components.

### 3.5 CIGB-300 stimulates cellular differentiation and cell cycle target genes

Myeloid hematopoietic differentiation is controlled by extrinsic cytokines and intrinsic transcription factors such as the macrophage colony-stimulating factor (M-CSF, encoded by CSF1) and EGR1. Both were studied by qPCR (**Figure S3**), showing an increase of the TF EGR1 in all conditions but mainly in HL-60 treated for 30min. CSF1 was significantly increased after 3h of HL-60 treatment with CIGB-300.

We also explored gene expression of CDKN1A/P21, with important regulator roles in G1 progression in cell cycle, by qPCR in both cell lines after the CIGB-300 treatment for 30 min, 2 h and 8 h. **Figure S4** shows the increase of CDKN1A mRNA levels in both cell lines; from 30 min in OCI-AML3 and after 2 h and 8 h of CIGB-300 treatment in HL-60 with higher magnitudes.

### 3.6 CIGB-300 elicits up-regulation of TNFA and NF-kB target genes

NF-kB pathway genes were shown to be significantly up-regulated by CIGB-300. To assess the robustness of the microarray analysis, we selected representative genes for validation by quantitative real-time RT-PCR (qPCR) and ELISA. Ten genes from the NF-kB signaling pathway, identified by the high-throughput analysis in HL-60 and OCI-AML3 cells treated with CIGB-300 in a temporal serial at 30 min and 3 h, were investigated by qPCR (**Figure 6**). The expression patterns obtained by PCR confirmed microarray results, in a cell-line and temporal manner, with higher absolute up-regulation fold changes values in relation to those in the microarray.

**Figure 6.**
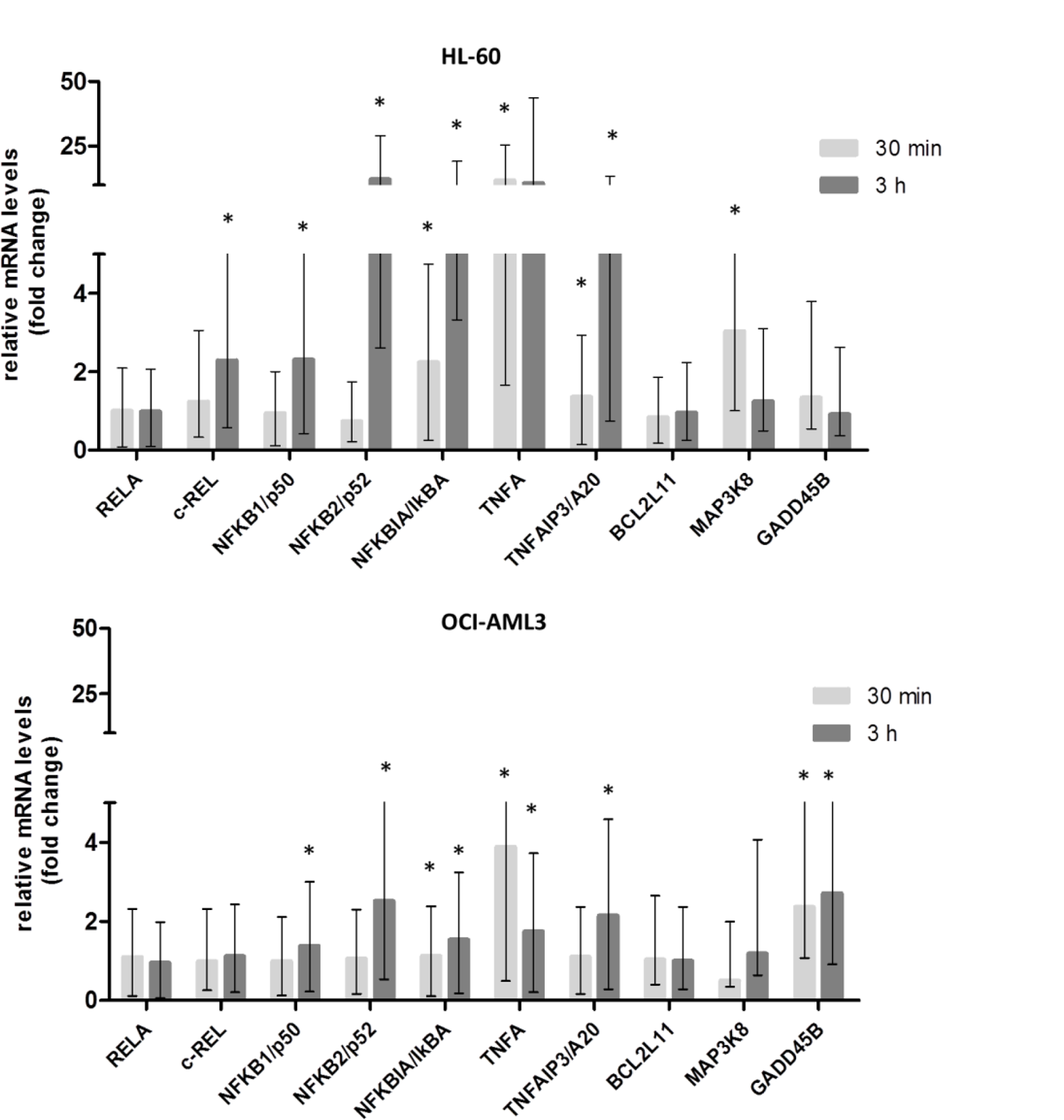
qPCR validation of selected genes of NF-kB signaling pathway modulated by CIGB-300. HL-60 and OCI-AML3 cells were treated with 40 µM of CIGB-300 for 30 min and 3 h. Histogram bars indicate relative mRNA levels ± standard deviation with respect to a time-matched untreated control, for two independent experiments analyzed in triplicate. All genes were normalized with ABL1, DDX5, and GAPDH genes. Asterisks represent statistically significant changes (p< 0.05) by REST 2009.

Target genes NFKBIA/IKBα, NFKB1/p50, NFKB2/p52, and TNFAIP3/A20 showed little or no over-expression early (30 minutes) but at 3 h they significantly increased expression in both cell lines. These increased levels were higher in the HL-60 cell line compared to OCI-AML3. c-REL/REL showed this same behavior in HL-60. In contrast, RELA gene levels were not significantly modulated by CIGB-300 in the microarray and qPCR analysis, neither the c-REL & RELA target BCL2L11. Moreover, c-REL targets MAP3K8 and GADD45B increased gene expressions either in HL-60 (30min) or in OCI-AML3 (30min and 3h), respectively.

In agreement with the qPCR result, a clear increase of the p50 binding activity after CIGB-300 treatment was confirmed by ELISA assay in both AML cell lines; although a higher level of p50 DNA binding was observed in OCI-AML3 cells compared to HL-60 (**Figure 7A**).

**Figure 7:**
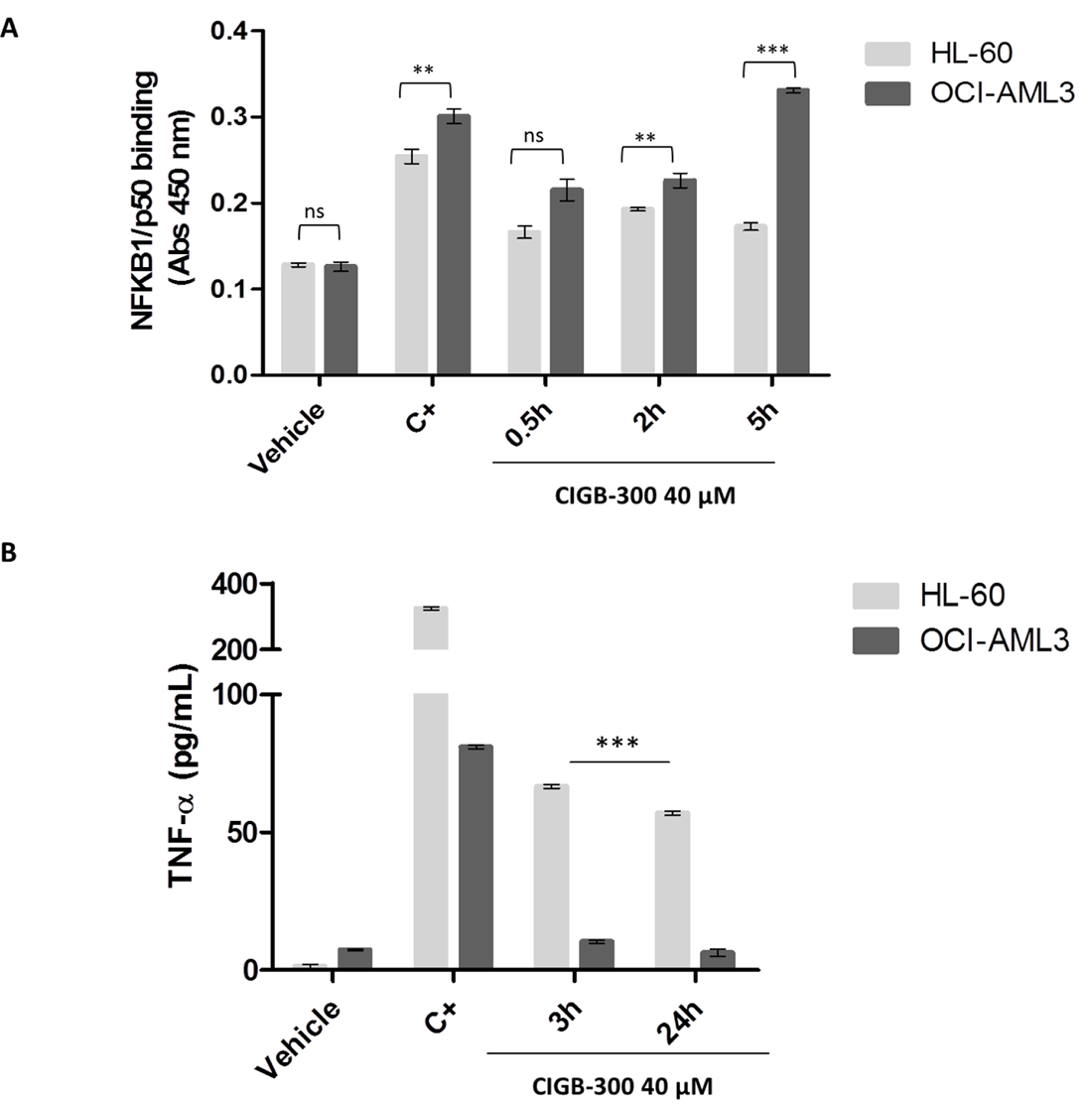
ELISA assays for p50 binding and TNF-α levels detection after CIGB-300 treatment of HL-60 and OCI-AML3 cells. (A) Cell extracts from HL-60 and OCI-AML3 cells were subjected to an ELISA Panomics kit (#EK110) to quantify p50 DNA binding activation after treatment with 40 µM of CIGB-300 at 0.5h, 2h, and 5h. NF-kB activation positive control (C+) was included. Histogram bars indicate Absorbance (Abs) 450nm mean± SD (standard deviation) for two independent experiments analyzed in triplicates. (B) Soluble TNF-α level was measured by ELISA after 3 h and 24 h of treatment with 40 µM of CIGB-300. PMA was employed as a positive control (C+) of TNF-α secretion. Histogram bars indicate pg/mL mean± SD for two independent experiments analyzed in triplicate. Statistically significant differences between conditions are represented as **p < 0.01 and *** p < 0.001 after one-way ANOVA followed by a Dunnett post-test.

qPCR results also evidenced TNFA(TNF-α) was up-regulated by CIGB-300 3.8-fold in OCI-AML3 and 11.6-fold in HL-60 as early as 30 minutes of drug exposure. This up-regulation was still found at 3h (1.7-fold for OCI-AML3 and 10.6-fold for HL-60). An ELISA assay to measure TNF-α secretion in AML cell lines following CIGB-300 treatment corroborated microarray and qPCR data showing a cell line-specific regulation with elevated TNF-α expression in HL-60 (66.7 pg/mL) compared to OCI-AML3 (10.3 pg/mL) at 3 h of treatment (**Figure 7B**). TNF-α induction by CIGB-300 tends to decrease at 24 h post-treatment, pointing to the time-dependent effect of this signaling pathway activation.

## 4. Discussion

AML is still a challenging disease in terms of effective therapy and minimal residual disease control [40], thus novel therapeutics have evolved in the last years, where CK2 targeting looks among the most promising ones [20, 41, 42]. Preclinical results have shown CIGB-300 peptide impacts on leukemic cell proliferation [21]. Preliminary clinical results show safety and first insights into the effect on AML patients [43].

*In vitro* models are often used to understand the cellular response, molecular mechanisms, and key pathways involved in the effect of a drug. Here, HL-60 (FAB M2 14% in AML; TP53/CDKN2A/NRASmut) and OCI-AML3 (FAB M4 20% in AML; DNMT3Amut, NPMc+) cell lines were chosen to cover an important part of AML patients with non-assigned therapy [in contrast to it is the case of FAB-M3] [44] [45]. Previous proteomic and phosphoproteomic analyses in both cell lines showed common but also specific ways CIGB-300 impacts cell apoptosis and/or the cell cycle [21, 24]. Gene expression profiles in both cell lines would also allow sensing what are general, or more specific mechanisms in relation to genetic backgrounds and differentiation states of leukemic cells. This study will give a new view at an earlier step of regulation. This is the first report of the gene expression profile modulated for CIGB-300 in HL-60 and OCI-AML3 cell lines using a microarray approach.

The Multidimensional Scaling and the ANOVA R^2^ of covariates analysis showed the main differences are governed by the cell line background and later, by treatment and time. Nevertheless, the heatmap picture showed a Cluster I comprising genes that are up-regulated in both cell lines although with differential magnitudes and kinetics. Interestingly, this cluster is composed of genes encoding transcription factors responding to stress stimulus as EGR1 and AP-1 components, pro-inflammatory cytokines such as IL1β and TNFα, and chemokines, including IL8. Most of them are part of the Top 100 up-regulated genes in the complete experiment, with a higher increase in HL-60. NFAT Transcription factor pathway was found enriched in both cell lines. Although the Calcineurin–NFAT pathway was described in T cells as acting as a master regulator of lymphocyte development and effector T-cell functions [46], it is also essential in myeloid lineages for their defense function against pathogens [47]. NFAT factors can cooperate with transcription factors such as AP-1 and NF-kB to modify immune responses. NFAT expression also increased in murine bone marrow cultures stimulated with M-CSF (encoded by CSF1) [48]. In normal physiology, M-CSF triggers hematopoietic stem cells to differentiate into macrophages/monocytes and in our experiment, CSF1 increased their gene expression after 3h of peptide treatment in HL-60. Leukemia-derived cells lines come from hematopietic precursors with potential to differentiate into immunocells. Ramírez et al (2017) profiled HL-60 promyelocytes differentiating into macrophages, neutrophils, monocytes, and monocyte-derived macrophages in a time series experiment from 3h to 168h [49]. Similarly, they found a rapid response at 3h with the expression of key transcription factors and lineage markers, including EGR1, EGR2, RELB, and NFKB2 in macrophage of both subtypes and NR4A1, NR4A2, NR4A3, EGR3 and FOSB in monocyte-derived macrophages. Heatmap of other differentially expressed transcriptional regulators showed the increase, as early as 3h, of AP-1 elements, REL/REL A, and inflammatory molecules such as IL1B or CCL2. This differentiation program is very well controlled over time with waves of expressions. NFAT pathway can also be stimulated via pattern recognition receptors (PRR) which increases the expression of IL2, −10, or −12. IL-10 signaling is enriched after 30min of treatment in OCI-AML3. These elements joined to the enrichment of regulation of hematopoiesis as biological process, along the experiment for the M2 cell line HL-60, and regulation of cell differentiation in both cell lines with different kinetics are indicating CIGB-300 possibly triggers differentiation in leukemic cells. Corroboration should be done measuring differentiation membrane markers in proper controlled experiments.

This transcriptomic analysis has particularly revealed the stimulation of cytokine and chemokine responses, as a quick molecular response to CIGB-300 and the involvement of an inflammatory response as part of the mechanism of this peptide. These CIGB-300 targeted cell lines are myeloid precursor in an intermediate state of differentiation that can evolve to become into effector myeloid cells and then their response could mimic that of an antigen-presenting cell. The presence of the cationic Tat penetrating peptide as part of the CIGB-300 sequence contributes to its rapid internalization within 3 min in 80 % of the cells [21]. Previous reports with other penetrating peptides have shown the increased uptake by antigen-presenting cells and the induction of a potent CD8^+^ T cell immunity. Using OVA as a vaccine antigen model, a potent antigen-specific immune response was induced with increases in IgG titer, splenocyte proliferation, secretion of cytokines IFN-γ, IL12, IL4, and IL10, immune memory function, and the activation and maturation of dendritic cells [50]. CIGB-300 has not elicited an antibody response after mice immunization (*unpublished results*), thus additional mechanisms where immune inflammatory response can contribute to change cells phenotypes, recruit effector immune cells nearby leukemic cells and then help to control cancer cell proliferation should be participating. This inflammatory response is higher in HL-60 after 3h of treatment, involving IL18 signaling. This pathway shares components with IL1 to activate NF-kB signaling, adhesion molecules, chemokines, IFN-γ, IL4/IL13 and Fas ligand [51]. Cell chemotaxis process is a process enriched in the HL-60 cell line while IL4/IL13 signaling was enriched at 3h in OCI-AML3. Heatmap Cluster II comprised genes participating in the positive regulation of leukocyte-mediated immunity, that only down-regulated expression in OCI-AML3 at a 3h time point. Evidently, both cell lines interpret initial inflammatory signaling in different ways and timing.

Higher number of regulated genes in HL-60 compared to OCI-AML3, mainly after 3h of treatment, indicates both models have their own ways to achieve the anti-proliferative state. Based on DEGs, iRegulon plugin predicted transcription factors with a NES score, and showed the early activation AP-1 components (JUN or JUND) and NF-kB family members together with EGR and other TF. The role of AP-1 and NF-kB family members is reinforced after 3h of peptide treatment only in HL-60, where JUNB, FOS, and REL are incorporated into the prediction. Networking analysis also showed a high degree of connections from several of these nodes. The vision of time dependence as well as their centrality to interact with other components of the functional net, including those genes participating in Apoptosis and Cell Cycle is better for HL-60 with 304 functional links. Although TFs are very well connected in the net for OCI-AML3, the ramification to the other genes is less evident showing that as few as 42 genes are functionally linked.

Singh et al (2011) showed AP-1 transcription factor family members’ c-Jun and JunB were transcriptionally activated in non-steroidal anti-inflammatory drugs treated AML cells, leading to the activation of GADD45A with Apoptosis induction [52]. CIGB-300 has been shown to have an impact on apoptosis in AML cells [21] and the process was found enriched in these cell lines after a proteomic experiment [24]. Extrinsic apoptotic plays an important role in the immune surveillance of transformed cells and it is activated by cytokine ligands binding (i.e., FasL, TNF, and TRAIL) to members of the TNFα receptor superfamily, also called death receptors (i.e., Fas, TNF, and TRAIL receptors), followed by the activation of an active caspase-8 and beyond effector caspases (caspase-3, −6 and −7) [53]. The pro-apoptotic effect of CIGB-300 in OCI-AML3 cells was supported in the proteomic experiment by the increase of BAK, FADD, caspase-7, and gasdermin-D levels [24]; while in HL-60, elements that counteract pro-apoptotic stimuli, probably to allow DNA damage repair under low levels of genotoxic stress, were over-expressed. In our experiment, FAS gene expression increased by 1.55 at 3h of treatment and TRAIL (encoded by TNFSF10) progressively increased along the experiment in HL-60. Preclinical models have shown recombinant TRAIL induces tumor regression with little toxicity to normal tissues [54]. In OCI-AML3, this apoptotic pathway showed no other changes that the increase in TRAIL receptor (encoded by TNFRSF10A) at 3h with a low FC of 1.27. In accordance, Positive regulation of cell death and Death Receptor signaling were shown as enriched Biological Processes in the Metascape tool at 3h in HL-60, including genes FAS, TNF, TNFRSF12A, TNFSF10, and also BCL2L11, encoding for the pro-apoptotic BIM in the intrinsic pathway [53].

NFAT members also participate in the transcription of several cytokines and signaling molecules with the activation or inhibition of Cell cycle [55]. The CDK inhibitor P21 is also regulated by this transcription factor. Here, we found CDKN1A (encoding P21) up-regulated after 3h of CIGB-300 treatment in both cell lines. In HL-60, a higher number of CDKN1A network interactions were found and a higher mRNA expression after 2h and 8h of CIGB-300 treatment was also demonstrated by qPCR. In contrast, OCI-AML3 showed the highest P21 increases at 30 min, although with lower values than in HL-60. Balusu et al. demonstrated NPM1 knockdown induced P53 and P21 and decreased the percentage of cells in S-phase of the cell cycle, thus this could be also the relation in this NPM1-mutated cell line [56]. The cell cycle related gene encoding GADD45B, targeted by P53 and c-REL, also showed differential regulation in both cell lines after qPCR validation. The modulation of P21 and GADD45B by CIGB-300 in both cell lines is another example of how the antiproliferative response could be achieved depending on cell background.

The induction of TNF response is evident in all groups of this experiment, with higher increases reached in HL-60. Four putative NFAT-binding sites, but also EGR1 and AP-1 binding sites, have been demonstrated in the TNF-α promoter, a gene that also increased. Systemic administration of TNFα in cancer therapy has been avoided because of its pro-inflammatory actions but TNF destroys tumor-associated blood vessels by apoptosis and improves vascular permeability to cytotoxic drugs. Thus a more controlled application displays advantages. Curnis et al (2000) showed that low doses of TNF improved penetration of doxorubicin in the treatment of melanoma and lymphoma [57]. Here, we showed increases in the gene encoding TNFα in both cell lines although increases were higher in the HL-60 cell line at gene and protein levels. Moreover, TNFα induction is transient as the protein expression is decreased 24h after peptide treatment compared to 3h. TNF receptor 1 (TNFR1) signaling is bifurcated into three different paths affecting cellular fate. In the absence of TRADD, ligand binding to TNFR1 recruits RIP1 producing reactive oxygen species (ROS), activating the JNK signaling cascade, and cells then die by apoptosis [58]. Incubation of AML cells with CIGB-300 peptide increased ROS production in HL-60 cells but not in OCI-AML3 cells. As a consequence, the HL-60 proteomic profile included ROS metabolic as an enriched process [24] and the connection between ROS de-regulation and CIGB-300-induced apoptosis was demonstrated. ROS can sustain JNK activity allowing TNFα to kill cells in which NF-kB is active [53]. In the other hand, autocrine binding of TNFα to TNFR2 up-regulates the anti-inflammatory cytokine IL10 in monocytes and clears TNF from the environment, serving as a balance for the pro- and anti-inflammatory actions of this cytokine [59]. As we mentioned before, IL10 signaling was enriched in OCI-AML3 at 30 min. This could be part of the explanation for the low TNFα detected in the supernatants of OCI-AML3 (10.3 pg/mL) compared to HL-60 (66.7 pg/mL) at 3h of treatment. The measure of TNFR1&TNFR2 membrane expression, as well as TNF secretion in a time course experiment, could clarify possible mechanisms and contributions in both models.

TNF signaling is also linked to NF-kB signaling. Although NF-kB signaling pathway has been mainly associated with a survival response in cancer to protect cells from apoptosis [60] different drugs targeting this cascade have been developed to control cancer cell proliferation [61]. Dual functions of NF-kB signaling appear to result from the ability of these transcription factors to either activate or repress transcription of genes depending on interaction with transcription co-activator or repressors and post-translational modifications [62, 63]. For instance, DNA-damaging chemotherapeutic agents can lead to NF-kB activation by initiating signals generated in the nucleus [64][65]. However, modulation of the NF-kB pathway and temporal dynamic depends on the cell type and the nature and amount of the agent [66, 67]. At present, the molecular characterization of the effect of CIGB-300 on the NF-kB signaling pathway is not fully elucidated. As a preliminary study we explored the gene expression of different components of canonical NF-kB signaling pathways by qPCR. In both AML-cell lines, we detected an over-expression of NFKBIA/IKBα, NFKB1/p50, NFKB2/p52, and TNFAIP3/A20 at 3h post-treatment although higher in HL-60. Interesting, only in HL-60, c-Rel/REL shows that same increase at 3h, as well as its target MAP3K8. In contrast, although we did not find a gene regulation for REL or REL A in OCI-AML3 an increase in their target GADD45B was accounted during the experiment setting. The activation of this cascade in both cell lines was then validated by qPCR with different magnitudes and paths.

CK2 protein kinase phosphorylates IkB (NF-kB inhibitor/NFKBIA gene) which promotes its degradation and releasing of NF-kB complexes to enter into the nucleus [68]. NF-kB p65/REL A subunit is also phosphorylated by CK2 in Ser529; both phosphorylation events implicate activation of this transcription factor [69]. CK2 inhibitors, CX-4945 and CIGB-300, have evidenced a clear inhibitory effect on the NF-kB signaling pathway [21, 70, 71]. Particularly, in two non-small-cell lung cancer models, the anti-proliferative effect was accompanied by the inhibition of the CK2-dependent canonical NF-kB pathway, with reduced REL A/p65 nuclear levels and conditionally reduced NF-kB transcriptional activity [71]. In the AML context, we demonstrated an up-regulation of some of the NF-kB target genes as well as TNF-α as one of these pathway inducer, following treatment with CIGB-300. This fact could be related to the activation of apoptosis as molecules from the mitochondrial (intrinsic) and death receptor (extrinsic) apoptotic pathways (ex. p53, Fas and FasL, TNFα, TRAIL, receptors DR4, DR5, DR6, pro-apoptotic Bcl-2 family members) are NF-kB transcriptional targets [53]. Certainly, a closer inspection of whether NF-kB signaling activation in AML cells triggers the apoptotic machinery induced by CIGB-300 or cell rescue signals upon cell exposure to the drug remain to be established.

In conclusion, we present the temporal gene profile in two cellular models for AML, HL-60 and OCI-AML3, treated with the peptide CIGB-300. An interesting finding at this early step of regulation is the increase of TF, cytokines and chemokines related to stress stimulus and inflammation since 30min of treatment which increases at 3h; TNF and NF-kB signaling are also stimulated at a higher proportion in HL-60. In this cell line, IL18 signaling is enriched in contrast to IL10 and IL4/IL13 signaling in OCI-AML3. Hematopoiesis and differentiation are also stimulated. Even when other experiments have shown the role of the peptide over apoptosis and cell cycle, this microarray profile gives new clues about the role of TNF and NF-kB signaling to accomplish those effects. The networking from several TF show the diversification to processes actors and the particular ways both cell lines achieve similar biological effects. Temporal gene expression shows that together with the anti-proliferative mechanism, CIGB-300 can stimulate immune responses by increasing immunomodulatory cytokines which, when locally secreted, may activate the immune system for tumor attack and blast cell control. This dual mechanism underscores this peptide as a promising AML therapeutic.

## 5. Author statements

Conceptualization, D.V.-B., Y.P. and S.E.P.; methodology, D.V.-B., A.C.R., M.R., A.R. and Y.P.; D.V.-B. and D.P followed the samples delivery and microarray experiment service; Formal analysis, D.V.-B., A.C.R., M.R., G.V.P., A.R., Y.P. and S.E.P.; Project administration, D.P. and S.E.P.; Writing—original draft preparation, D.V.-B. and A.C.R.; Writing—review and editing, M.R., G.V.P., Y.P. and S.E.P. All authors have read and agreed to the published version of the manuscript.

## Institutional Review Board Statement

Not applicable.

## Informed Consent Statement

Not applicable.

## Data Availability Statement

Additional information would be available from the corresponding authors upon reasonable request

## Acknowledgments

We would like to thank to all the parts involved at the CIGB for microarray service and sample delivery to McGill University and Génome Québec Innovation Centre (Montréal, Québec, Canada). We thank to MSc. Jamilet Miranda (Bioinformatic Department, CIGB) for her initial contribution to gene expressions explorations in AML cell line treated with CIGB-300.

---

**Figure S1.**
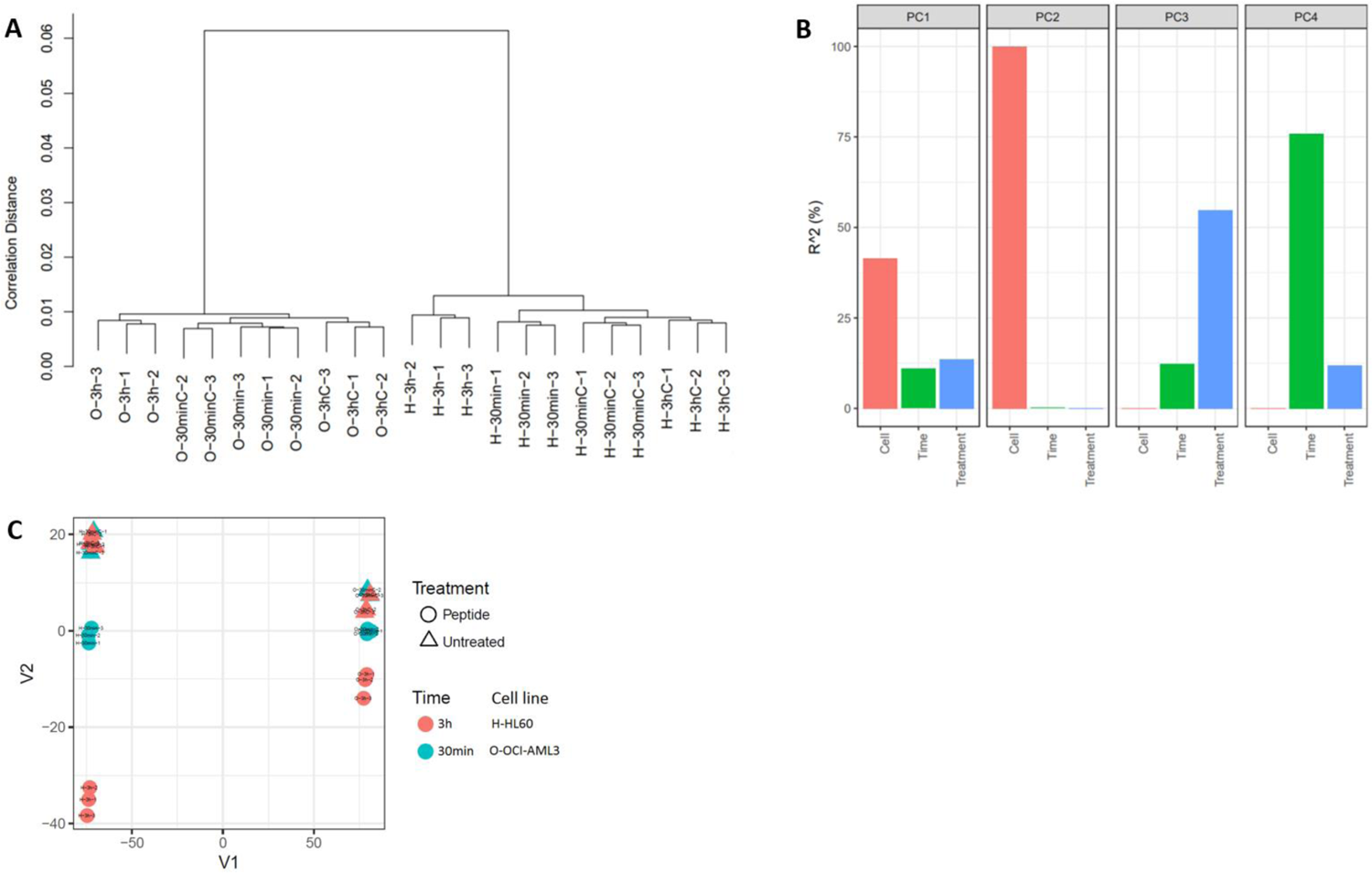
Diagnostic plots in control cells or CIGB-300 treated OCI-AML3 (O) and HL-60 (H) samples for 3h and 30min. (A) One dimensional hierarchical clustering of all samples (replicates 1, 2, 3), (B) ANOVA R-squared (^2) of covariates (Cell, Time and Treatment) in % (C), Multidimensional Scaling (MDS) of filtered data.

**Figure S2.**
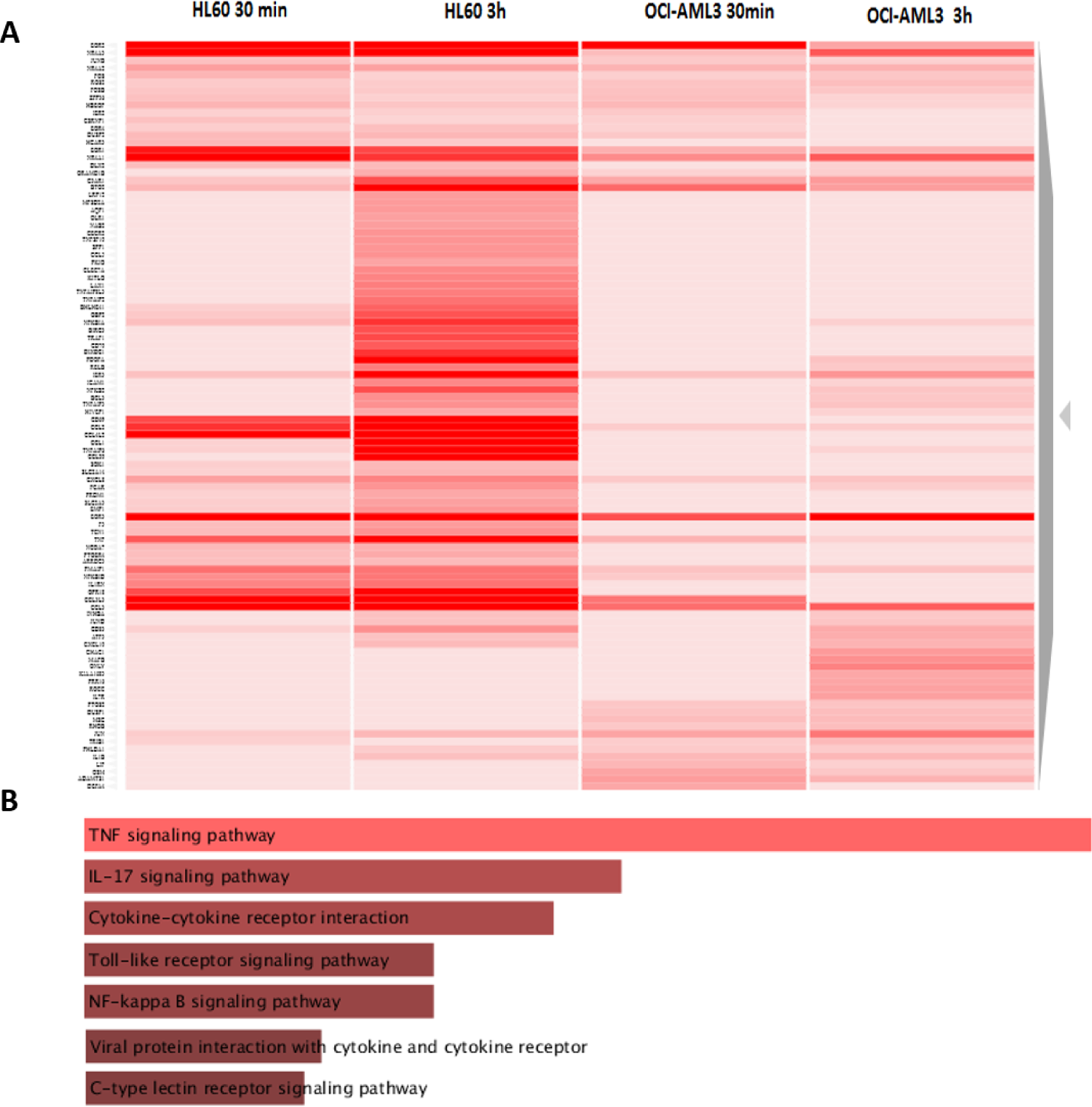
Top 100 differentially up-regulated genes by CIGB-300. (A) Unsupervised Heatmap from the Top 100 most differentially up-expressed genes in HL-60 and OCI-AML3. (B) Pathways analysis in Top 100 up-regulated genes using Enrichr.

**Figure S3.**
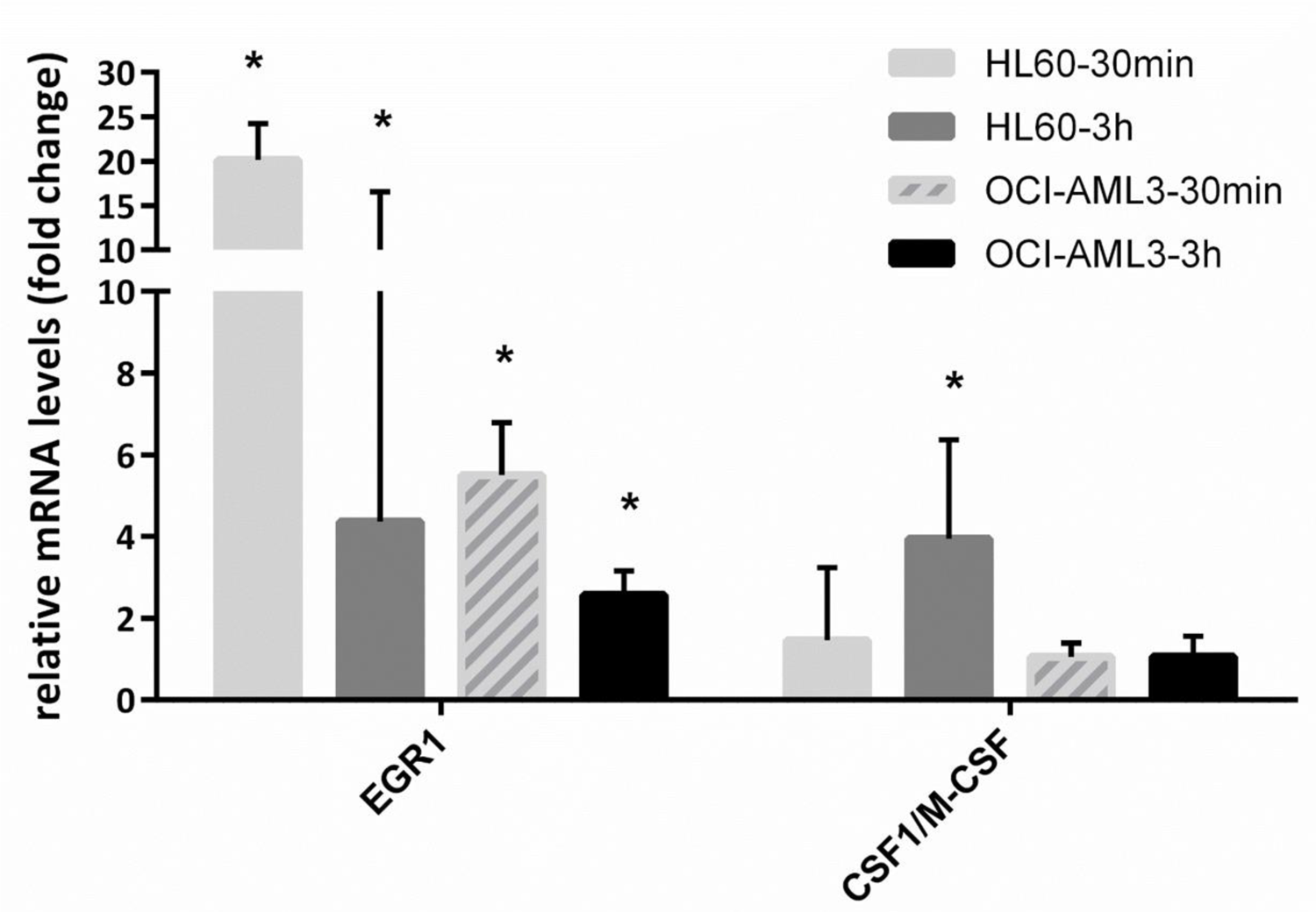
qPCR validation of genes related to differentiation modulated by CIGB-300. HL-60 and OCI-AML3 cells were treated with 40 µM of CIGB-300 for 30 min and 3 h. Histogram bars indicate relative mRNA levels ± standard deviation with respect to a time-matched untreated control, for two independent experiments analyzed in triplicate. All genes were normalized with ABL1, DDX5, and GAPDH genes. Asterisks represent statistically significant changes (p< 0.05) by REST 2009.

**Figure S4.**
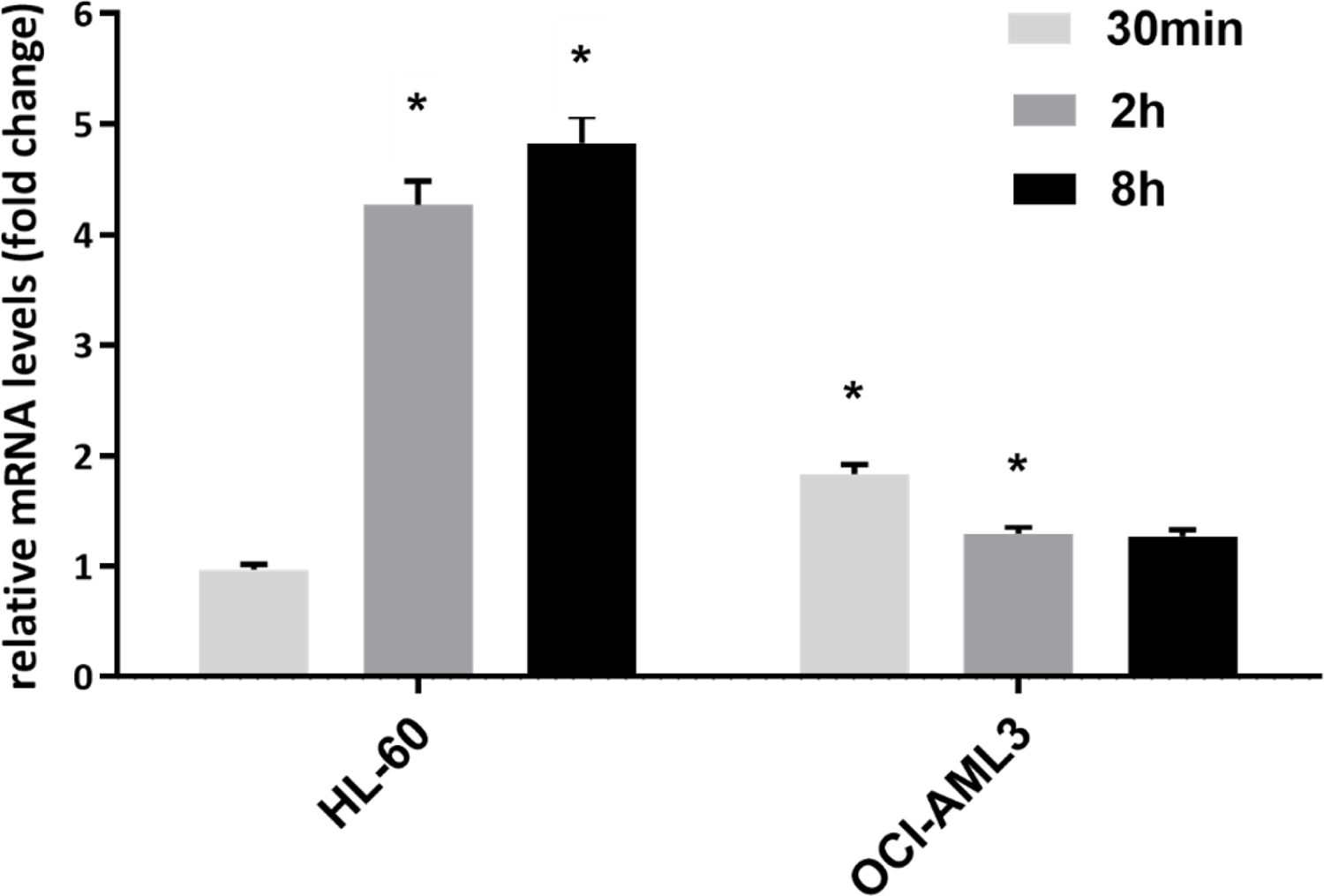
qPCR validation of the cell cycle related gene CDKN1A/P21 gene modulated by CIGB-300. HL-60 and OCI-AML3 cells were treated with 40 µM of CIGB-300 for 30 min, 2 h and 8 h. Histogram bars indicate relative mRNA levels ± standard deviation with respect to a time-matched untreated control, for two independent experiments analyzed in triplicate. All genes were normalized with ABL1, DDX5, and GAPDH genes. Asterisks represent statistically significant changes (p< 0.05) by REST 2009.

